# Sensitization of cutaneous primary afferents in bone cancer revealed by *in vivo* calcium imaging

**DOI:** 10.1101/2020.09.01.275099

**Authors:** Larissa de Clauser, Ana P. Luiz, Sonia Santana-Varela, John N. Wood, Shafaq Sikandar

## Abstract

Cancer-induced bone pain (CIBP) is a complex condition comprising components of inflammatory and neuropathic processes, but changes in the physiological response profiles of bone-innervating afferents remain poorly understood. We used a combination of retrograde labelling and *in vivo* calcium imaging of bone marrow-innervating DRG neurons to determine the contribution of these cells in the establishment and maintenance of CIBP. We found a majority of femoral bone afferent cell bodies in L3 DRG that also express the sodium channel subtype Na_v_1.8 - a marker of nociceptive neurons- and lack expression of parvalbumin - a marker for proprioceptive primary afferents. Surprisingly, the response properties of bone marrow afferents to both increased intraosseous pressure and acid were unchanged by the presence of cancer. On the other hand, we found increased excitability and polymodality of cutaneous afferents innervating the ipsilateral paw in cancer bearing animals, as well as a behavioral phenotype that suggests changes at the level of the DRG contribute to secondary hypersensitivity.

## 1 Introduction

Cancer patients experience pain throughout disease progression and even after treatment, with a prevalence of 60% during metastatic disease (Van Den Beuken-Van Everdingen *et al.*, 2016). A large number of solid tumours develop aggressive metastases to secondary sites, most notably to bone tissue (Coleman, 2000; Coleman *et al.*, 2006), and 38% of all cancer pain can be linked to nociceptor activation in the bone (Grond *et al.*, 1996). Cancer-induced bone pain (CIBP) produces intense episodes of breakthrough pain (Deandrea *et al.*, 2014; Scarpi *et al.*, 2014) that is unresponsive to conventional treatment (Mercadante *et al.*, 2004). Despite the implementation of WHO guidelines for pain management, analgesia remains inadequate in at least 22% and up to 100% of cancer patients (Carlson, 2016).

CIBP is a unique and complex condition involving inflammation, neuropathy and ischemia (Mantyh, 2013; Falk & Dickenson, 2014). The cancer and tumour-associated stromal cells, including fibroblasts, endothelial cells, lymphocytes, and many bone marrow-derived cells contribute to the inflammatory component of CIBP and are known sensitizing agents of sensory neurons (Julius & Basbaum, 2001; Joyce & Pollard, 2009). Nerve growth factor (NGF) also drives peripheral inflammation and sprouting of bone innervating sensory and sympathetic fibers, contributing to CIBP (J. M. Jimenez-Andrade *et al.*, 2010; Mantyh *et al.*, 2010; Bloom *et al.*, 2011). In accordance with this finding, immunohistochemical studies have revealed that over three-fourths of bone marrow afferents express the NGF receptor tropomyosin receptor kinase A (TrkA) (Juan Miguel Jimenez-Andrade *et al.*, 2010; Castañeda-Corral *et al.*, 2011).

Besides inflammatory and neuropathic processes in the bone microenvironment, cancer-induced bone pain is thought to be associated with increased intraosseous pressure (Sottnik *et al.*, 2015). This is mechanistically similar to intraosseous engorgement syndrome that leads to sensitization of primary afferents (Mantyh, 2014; Ivanusic, 2017). In rats, inflation of an intrafemorally implanted balloon produces nocifensive responses (Ishida *et al.*, 2016) and electrophysiological recordings demonstrate that tibial afferents are activated by intraosseous pressure stimuli produced by injection of different volumes of saline (Nencini & Ivanusic, 2017). Moreover, tibial afferents can be sensitized by inflammatory mediators such as capsaicin (Nencini & Ivanusic, 2017), NGF (Nencini *et al.*, 2017), carrageenan (Nencini, Thai & Ivanusic, 2018), and GDNF family ligands (Nencini *et al.*, 2018). Increased activity of bone resorbing osteoclasts (Neri & Supuran, 2011; Yoneda, Hiasa, Nagata, Okui & F.A. White, 2015) and a shift to aerobic glycolysis in cancer cells (Simon & Keith, 2008) leads to local acidosis within the hypoxic bone microenvironment. These events can also sensitize bone afferent neurons through transient receptor potential vanilloid type 1 ion channels (TRPV1) and acid-sensing ion channels (ASIC) to maintain CIBP (Yoneda *et al.*, 2011; Wakabayashi *et al.*, 2018). These findings support the notion that bone marrow afferents are a heterogenous population of sensory neurons that contribute to nociceptive processing.

Vast changes in bone homeostasis, combined with structural and neurochemical reorganization of sensory and sympathetic nerve fibers in the bone, highlight the importance of peripheral mechanisms underlying CIBP. Clinical evidence suggests peripheral input is required for the maintenance of metastatic bone cancer pain (Klepstad *et al.*, 2015) and peripheral nerve block reverses tactile hypersensitivity and impaired limb use in a rat model of CIBP (Remeniuk *et al.*, 2015). Here, we use a combination of retrograde and genetic labeling with *in vivo* calcium imaging of primary afferents at the level of the soma in order to investigate whether bone marrow innervating neurons play a role in the establishment and maintenance of pain in animals with metastatic bone cancer. In this study we (1) identified the molecular profile, size and distribution of sensory neurons innervating the femoral bone marrow; (2) defined the role of single bone marrow afferents in nociceptive signaling in a murine model of bone cancer; and (3) determined the contribution of DRG sensory neurons to secondary hypersensitivity observed at distal sites to the tumour bearing bone.

## 2 Results

### 2.1 Molecular characterization of bone afferents

We performed retrograde tracing of bone marrow afferents in different tdTomato reporter lines to produce a molecular expression profile of sensory neurons innervating the bone (Figure 2 and Table 1). We first determined the rostrocaudal distribution of bone marrow afferents. We found the majority of cell bodies of retrogradely labelled neurons in L3 DRG (34.73±10.13%), followed by L4 (32.98±9.46%) (n=3 animals) (Figure 2A). To account for differences in total number of neurons within each lumbar DRG, we determined the proportion of bone marrow afferent neurons within the population of all DRG neurons expressing Na_v_1.8 (using tdTomato reporter fluorescence). The highest percentage of bone afferents were found in L3 DRG, although L3 and L4 proportions did not significantly differ (Figure 2B). Based on these findings we focused on L3 DRG to determine the molecular identity of bone afferents. A total of 287 retrogradely labelled bone marrow afferents were counted in L3 DRG from 13 animals (average of 22.08±3.50 per mouse). Neuronal size was defined as small (<300 μm^2^), medium (300-700 μm^2^) and large (>700 μm^2^) as previously described (Ruscheweyh *et al.*, 2007). The majority of bone marrow afferents are medium sized (48%), followed by large sized (37%), and small neurons (15%) (Figure 2C). A substantial proportion of L3 bone marrow afferents (73.80±5.81%, n=5 animals) express Na_v_1.8, and most of these neurons show TrkA immunoreactivity (94.05±1.68%, n=2 animals) (Figure 2D). One-fifth (20.38±6.00%, n=3 animals) of bone marrow afferents express Tmem233, and the vast majority of these neurons show TrkA immunoreactivity (93.75±3.61%, n=3 animals) (Figure 2E). One single bone afferent was found to express Pvalb (0.71±0.64%, n=5 animals) (Figure 2F, Table 1).

**Figure 1.**
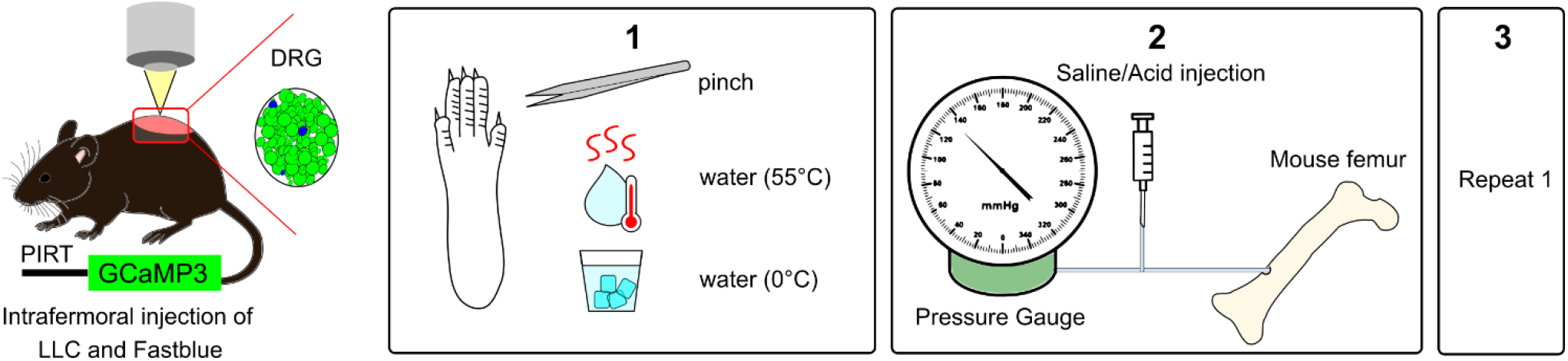
Experimental setup for *in vivo* calcium imaging of cutaneous and bone marrow afferents. Adult mice expressing GCaMP3 under the control of the promoter Pirt were injected with Lewis Lung carcinoma (LLC) cells or vehicle and Fast Blue retrograde tracer into the distal femur head. (1) To measure secondary hypersensitivity, noxious pinch, hot water (55°C) and ice-cold water (0°C) were applied to the plantar surface of the affected paw. (2) To activate bone afferents 10μl solution were delivered to the femoral marrow through a syringe and pressure was recorded. (3) Step 1 was repeated to determine if activation of bone afferents resulted in changes in excitability of distal cutaneous afferents.

**Figure 2.**
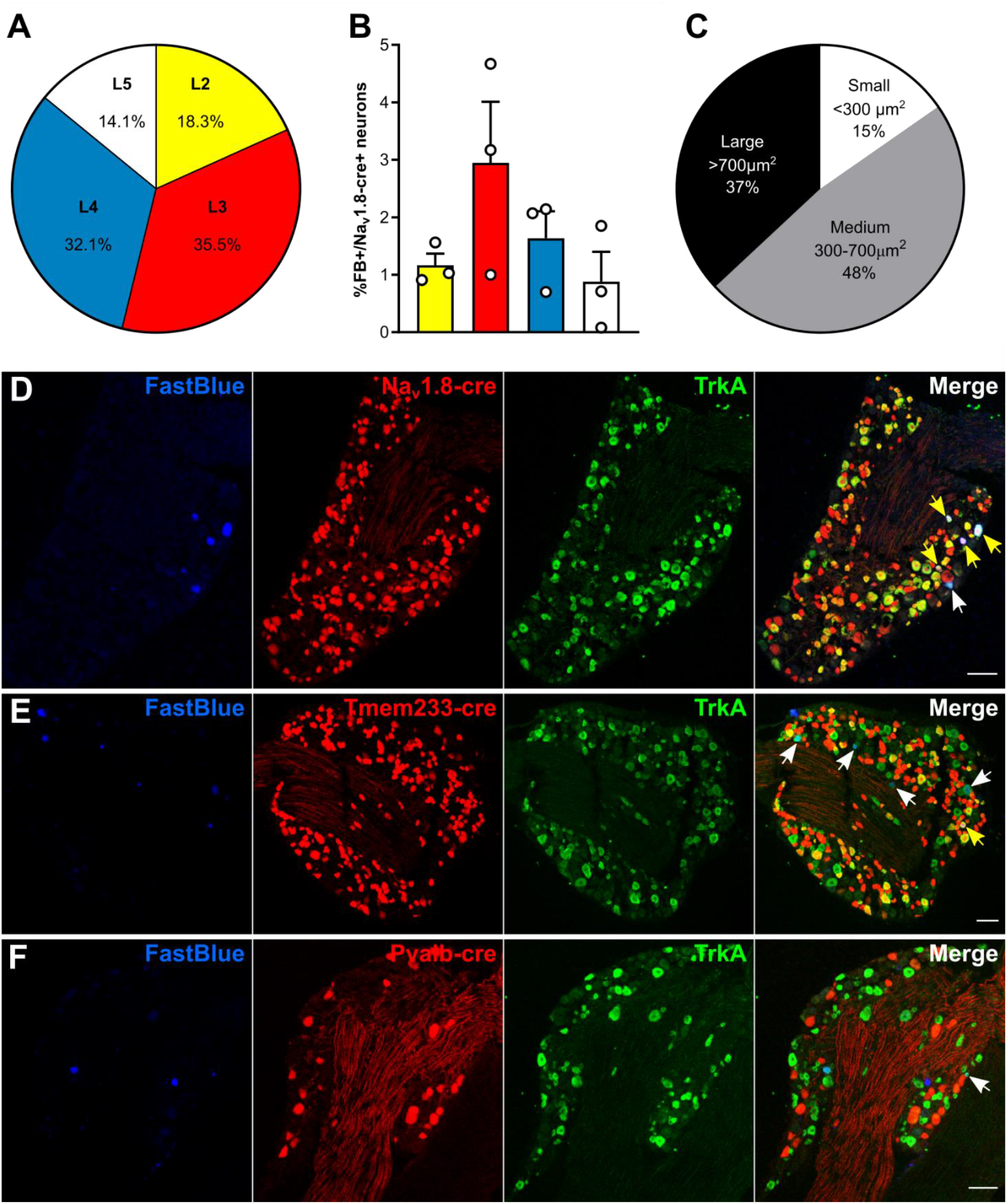
Bone marrow afferents are medium-to large sized neurons expressing the nociceptive marker Na_v_1.8 and lacking expression of proprioceptive marker parvalbumin. (A) Rostrocaudal distribution of all FB+ retrogradely labelled bone marrow afferents within the analyzed ipsilateral lumbar DRG (L2-L5). (B) Proportion of FB+ neurons within the Na_v_1.8-cre expressing population of DRG neurons throughout lumbar L2-L5. (C) Proportion of bone marrow afferents based on soma size. (D-E) Representative images of lumbar 3 DRG immunostained for TrkA (green) and Fast Blue retrogradely traced (blue) femoral bone marrow afferents in combination with tdTomato expressing neurons (red) driven by (D) Na_v_1.8-cre, (E) Tmem233-cre, and (F) Pvalb-cre. White arrows indicate double positive Fast Blue/TrkA neurons, yellow arrows indicate triple positive neurons. Scalebar = 100 μm.

**Table 1.**
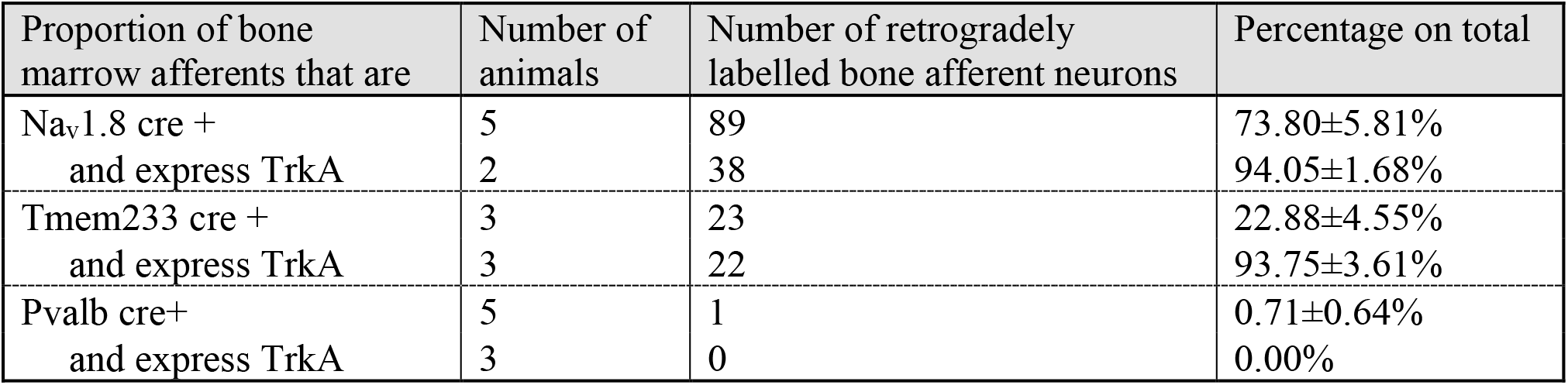
Expression of neuronal markers Na_v_1.8, Tmem233 and Pvalb in bone marrow afferent neurons.

To determine if bone afferents could be defined based on size within each subpopulation of sensory neurons, we compared the average sizes within each population. The mean size of the Na_v_1.8+ DRG population is significantly lower than the mean size of all bone afferents and Na_v_1.8-cre expressing bone afferents (Kolmogorov-Smirnov test: ****p<0.0001, Figure S 1A). Likewise, the mean size of the Tmem233+ population is smaller than the mean size of all retrogradely labelled bone afferents (Figure S 1B; Kolmogorov-Smirnov test: ****p<0.0001) and Tmem233+ bone afferents (Kolmogorov-Smirnov test: **p=0.0019) respectively. Size distribution of Pvalb+ neurons and bone afferents did not significantly differ (Figure S1 C; Kolmogorov-Smirnov test: p=0.0940).

### 2.2 GCaMP3-expressing mice show a moderate pain phenotype in a model of CIBP

The endpoint of the study was defined as animals showing clear signs of limping (Limb use score = 2 or 1), when cancer-bearing mice already show changes in gene expression at the level of the DRG (Bangash *et al.*, 2018). Cancer bearing mice started to limp between day 10 - 18 post-surgery, as outlined in the survival curve (Figure 3A) (Log-rank test, ****p<0.0001). Static weight bearing on the affected limb was markedly reduced at the endpoint in cancer animals compared to both baseline and their sham counterparts (Figure 3B) (two-way ANOVA with Bonferroni post-hoc, ****p<0.0001). Secondary mechanical hypersensitivity was assessed with von Frey filaments, and we found a significant difference in 50% withdrawal thresholds at the endpoint in cancer animals compared to baseline (two-way ANOVA with Bonferroni post-hoc, ***p=0.0002) and to sham animals (two-way ANOVA with Bonferroni post-hoc, ** p=0.0092) (Figure 3C). Primary hypersensitivity was assessed using non-noxious palpation of the distal femur head, which produced a marked increase in nocifensive responses (including guarding, flinching, and licking) in cancer animals compared to baseline (two-way ANOVA with Bonferroni post-hoc, *p=0.0378) and compared to sham animals at the endpoint (two-way ANOVA with Bonferroni post-hoc, *p=0.0159) (Figure 3D). On the other hand, pain pressure thresholds of the paw ipsilateral to the affected femur were comparable in sham and cancer bearing animals at baseline and endpoint (Figure 3E). Similarly, thermal pain thresholds of cancer animals measured using the hot-plate test at 50°C also indicated no differences between groups (Figure 3F).

**Figure 3.**
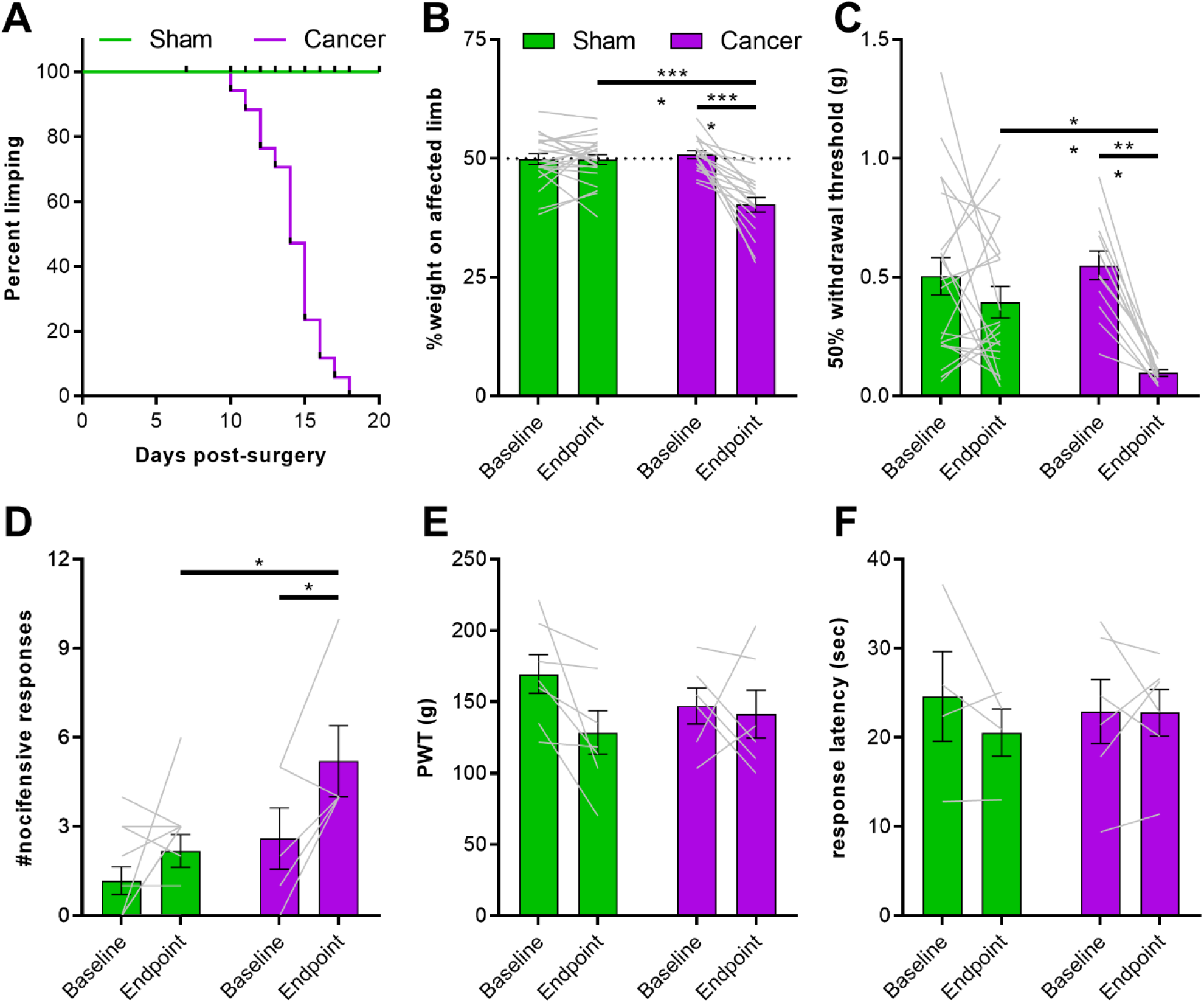
Pain behaviour in Pirt-GCaMP3-expressing mice with CIBP. (A) Survival curve after surgery for sham (green line, n=22) and cancer animals (purple line, n=17) with endpoint defined as clear limping on the affected limb. Black bars indicate individual dropouts. (B) Ongoing pain was measured by percentage weight bearing on the affected limb. (C) Mechanical withdrawal threshold to von Frey filaments in cancer (n=12) and sham animals (n=21). (D) Number of nocifensive responses (guarding, licking, flinching) observed during the 2min period after palpation of the distal femur head in cancer bearing (n=5) and sham (n=11) mice. (E) Mechanical withdrawal thresholds to application of the Randall-Selitto apparatus to the paw in cancer (n=6) and sham animals (n=7). (F). Thermosensation measured by response threshold to 50°C hot-plate in cancer (n=6) and sham mice (n=4).

### 2.3 Properties of cutaneous afferents in CIBP animals

To investigate the role of cutaneous afferents in secondary hyperalgesia of the hindpaw, a number of different stimuli were applied to the glabrous skin of the plantar surface for 3 sec at 30 sec intervals during *in vivo* calcium imaging. First, tweezers were used to apply pressure across the dermatome covering L3-L4, followed by hot water at 55°C, and ice-cold water. Cancer bearing animals presented a significantly larger proportion of neurons sensitive to pinch compared to sham animals, but a lower proportion of heat sensitive neurons (Figure 4A) (two-way ANOVA with Bonferroni post-hoc, ****p<0.0001). This shift may depend on increased polymodality in response to tissue injury or inflammation (Emery *et al.*, 2016). Polymodality was significantly increased in cancer bearing animals from 2.56% to 5.56% (Figure 4B) (Welch’s t-test, *p=0.0463). As *in vivo* imaging allows for the detection of cross-activation of sensory neurons, we investigated whether this mechanism contributes to the observed shift by determining the coupling response of DRG neurons (Kim *et al.*, 2016). The percentage of coupled responses within all mechanosensitive neurons, although higher in cancer bearing animals, did not significantly differ from sham mice (6.59±1.73% vs. 2.42±1.33% respectively; Welch’s t-test, p=0.0717). On the other hand, cross-activation of heat sensitive neurons was observed in very few cases both in sham and cancer animals (Figure 4C) (3.54±1.27% and 1.60±0.92% respectively; Welch’s t-test, p=0.1909). Changes in fluorescence of GCaMP-expressing DRG neurons during stimulus application (compared to baseline unevoked fluorescence) serves as a surrogate for the strength of the calcium transient. We found that the maximum response intensity of both mechanosensitive (Figure 4D) (1.32±0.07 in sham vs. 1.62±0.08 in cancer, Welch’s t-test, **p=0.0077) and heat sensitive (Figure 4E) (1.31±0.04 in sham vs. 1.62±0.07 in cancer, Welch’s t-test, ***p=0.0005) L3 cutaneous afferents were increased in cancer bearing compared to sham animals. To determine if previously silent nociceptors are recruited in response to noxious pinch, we compared size distributions. 170 mechanosensitive neurons in sham animals and 147 mechanosensitive neurons in cancer animals showed the same size distribution (Figure 4F) (Kolmogorov-Smirnov test, p=0.8275). Likewise, 366 heat sensitive neurons in sham and 147 in cancer animals did not show any difference in size distribution (Figure 4G) (Kolmogorov-Smirnov test, p=0.6651).

**Figure 4.**
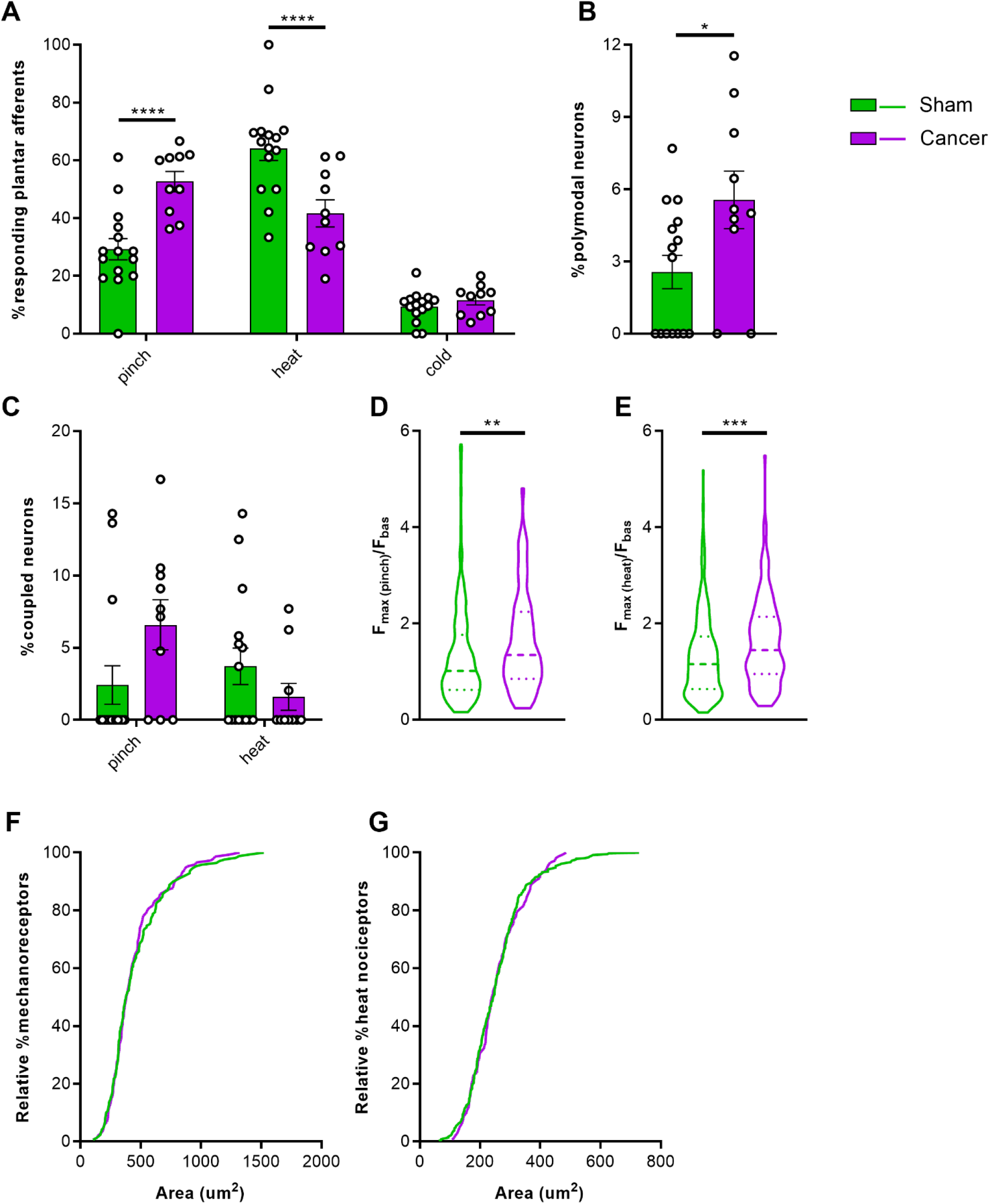
Response properties of cutaneous afferents of the hindpaw. (A) Percentage of total neurons responding to pinch, heat and cold in cancer (purple, n=10) and sham animals (green, n=15). (B) Percentage of polymodal cutaneous afferents in L3 DRG of cancer and sham mice. (C) Percentage of coupled responses over total responses of the same stimulus modality for pinch and heat in sham and cancer-bearing mice. (D-E) Response intensity of mechanosensitive (D) and heat sensitive (E) cutaneous afferents expressed as maximum fluorescence intensity during stimulus application vs. baseline. (F-G) Size distribution of mechanosensitive (n=147 neurons in cancer, n=170 neurons in sham) (F) and heat sensitive (n=147 in cancer, n=366 in sham) cutaneous afferents (G).

### 2.4 Activation of femoral bone marrow afferents

To investigate the contribution of intraosseous pressure and the acidic microenvironment in CIBP, we used intrafemoral injections of saline and citric acid, respectively, during *in vivo* DRG imaging of sham and cancer animals. In a subset of mice, we used a pump-controlled syringe connected to a pressure gauge to deliver 10 μl saline into the femoral marrow of sham and cancer bearing animals to serve as an indirect measure of intraosseous pressure (Figure 1). This initial pressure prior to injection of saline did not significantly differ between cancer and sham animals (125.60±18.96 and 86.67±13.33 mmHg respectively, Welch’s t-test, p=0.1440) (Figure 5A). The time required for delivering 10 μl saline in cancer animals (52.2±4.41 sec) was nearly threefold of that observed in sham animals (12.67±6.22 sec) (Welch’s t-test, **p=0.0020) (Figure 5B). To visualize bone marrow afferents during *in vivo* calcium imaging, mice were injected with Fast Blue retrograde tracer during tumour infiltration. Bone afferents were deemed to be activated (or ‘responders’) if a calcium transient was detected either during or after intraosseous injection (representative frames and traces taken from a recording are shown Figure 5C and 5D, respectively). We found that only a small proportion (8.5%) of Fast Blue retrogradely labelled neurons responded to intraosseous stimulation. However, 15% of all DRG neurons that were activated by intraosseous injection were also retrogradely labelled with Fast Blue (Figure 5E). The percentage of bone marrow afferents responding to stimulation of the paw following intrafemoral saline or acid injection did not significantly vary between cancer and sham animals (one-way ANOVA, p=0.6144) (Figure 6A). The response intensity of bone afferents also appeared to be unaffected by the presence of cancer (one-way ANOVA, p=0.4152) (Figure 6B). Although size distribution of responding bone afferents did not significantly differ between cancer and sham animals (one-way ANOVA, p=0.1762), a shift to the right was apparent in cancer bearing animals, suggesting that more medium-to-large sized bone afferents are recruited in nociceptive processing in CIBP (Figure 6C).

**Figure 5.**
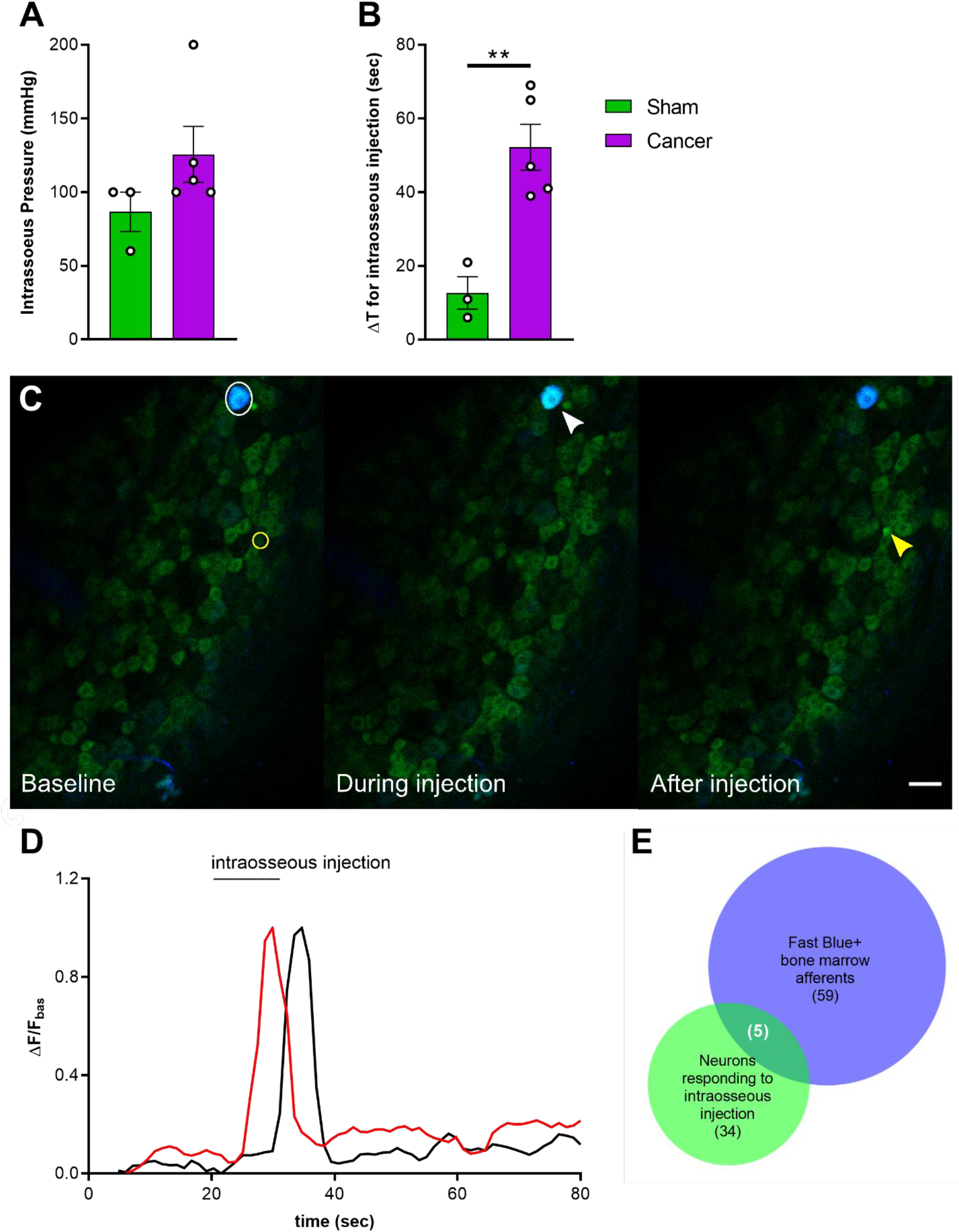
Activation of femoral bone marrow afferents through intraosseous injection. (A) Intraosseous injection pressure at start of solution flow in cancer (purple, n=5) and sham animals (green, n=3). (B) Time elapsed from start of solution flow till delivery of 10μl into the femoral bone marrow. (C) Representative images taken from a recording and showing the response of one Fast Blue labelled bone afferent (white arrow) responding during intraosseous injection and one unlabeled responding after (yellow arrow). Scalebar = 50 μm. (D) Example traces showing ΔF/F_basal_ of bone afferents responding during (red trace) or shortly after (black trace) injection into the mouse bone marrow. Both types of responders were included in the analysis. (E) Venn diagram showing the overlap between Fast Blue+ cells (blue) and neurons responding to bone marrow injection (green) (data from animals with specific Fast Blue labelling, n=6).

### 2.5 Response of plantar cutaneous afferents is unchanged after intraosseous stimulation

To investigate if acute stimulation of the bone marrow drives cross-activation of bone and cutaneous DRG neurons, the paw was stimulated again following injection of saline or acid into the femur. We found that the proportion of neurons responding to each modality was unchanged (Figure 6D) (two-way ANOVA within each group for each stimulus; cancer saline: p=0.0815 for pinch, p=0.1269 for heat; for all other groups and conditions: p>0.9999).

**Figure 6.**
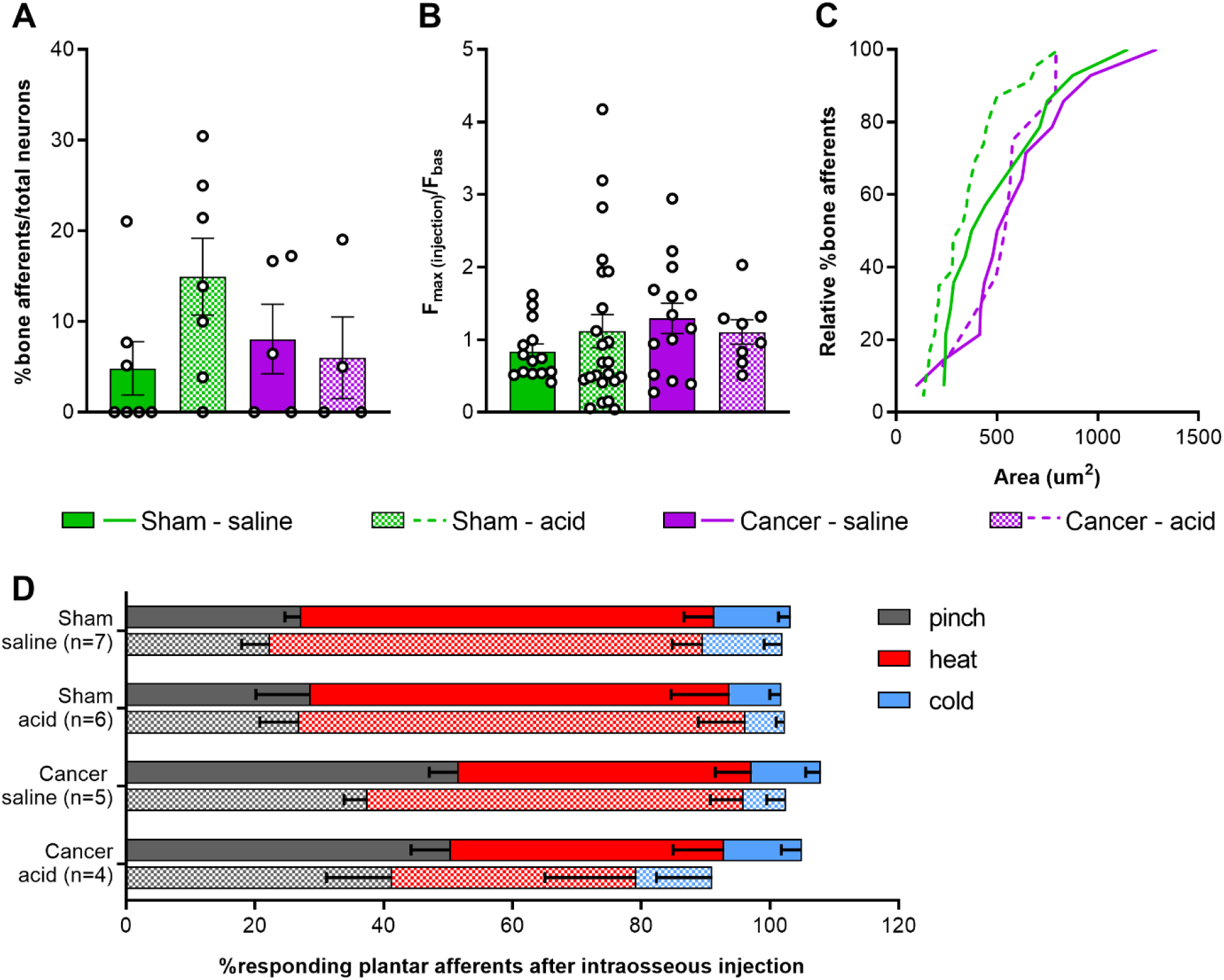
Response properties of femoral bone marrow afferents. (A) Percentage of bone afferents on total number of neurons responding to plantar stimulation in sham (green) and cancer animals (purple) receiving intraosseous injection of saline (solid fill, n=7 for sham, n=5 for cancer) or acid (pattern fill, n=7 for sham, n=4 for cancer). (B) Maximum fluorescence intensity of bone marrow afferents responding to intraosseous injection (number of neurons: n=14 for sham saline, n=14 for cancer saline, n=18 for sham acid, n=8 for cancer acid). (C) Size distribution of bone afferents responding to saline (full line) or acid (stitched line) in cancer and sham animals. (D) Percentage of responding cutaneous afferents before (solid fill boxes, pinch=grey, heat=red, blue=cold) and after (pattern fill boxes) stimulation of bone marrow afferents.

## 3 Discussion

### 3.1 Femoral bone marrow afferents are largely nociceptive and not proprioceptive

We first analysed the rostrocaudal distribution of mouse femoral bone afferent along lumbar DRG using retrograde labelling with Fast Blue. Consistent with sciatic and femoral nerve anatomy in the mouse (Rigaud *et al.*, 2008), we found the majority of somata of retrogradely labelled bone afferents in L3 DRG. Similar to previous studies in the rat, we also observed a range of soma sizes of bone afferents (Ivanusic, 2009; Nencini & Ivanusic, 2017), with most neurons classified as large- and medium-sized. Using three different reporter lines we identified the nociceptive neuronal marker Na_v_1.8+ to be expressed in over ¾ of bone marrow afferent neurons. This population largely overlapped with TrkA-expressing neurons, a known marker for these cells (Juan Miguel Jimenez-Andrade *et al.*, 2010; Castañeda-Corral *et al.*, 2011; Nencini *et al.*, 2017). Interestingly, ablation of Na_v_1.8-expressig neurons using Cre-mediated diphtheria toxin expression neither prevents nor attenuates the pain phenotype and bone degradation induced by intra-femoral injection of cancer cells (Minett *et al.*, 2014). This suggests that non-Na_v_1.8-expressing bone afferents, although a smaller population of DRG cells, are sufficient to induce physiological and nociceptive changes leading to the development of CIBP. Tmem233, a gene expressed within a subset of Na_v_1.8+ positive neurons, was present in 1/5^th^ of retrogradely labelled bone afferents, where it also largely overlapped with TrkA immunoreactivity. In addition, we observed an almost complete absence of Fast Blue retrogradely labelled cells in the Pvalb-cre+ neuronal subpopulation, indicating bone afferents do not play a role in proprioception. Single cell RNA sequencing data also demonstrates that the Pvalb-expressing population of mouse lumbar DRG neurons is distinct to the Na_v_1.8-expressing population of nociceptors (Usoskin *et al.*, 2015; Zeisel *et al.*, 2018)

### 3.2 Cancer bearing mice with a moderate reduction of limb use develop secondary mechanical hypersensitivity

Cancer bearing mice presented clear signs of limping, a reduction in weight bearing on the affected limb, and increased nocifensive responses to non-noxious palpation of the distal femur head, as has been previously reported (Luger *et al.*, 2002; Sabino *et al.*, 2003). Withdrawal thresholds to von Frey filaments applied to the ipsilateral paw were substantially reduced, while withdrawal to the application of the Randall-Selitto pressure clamp to the ipsilateral paw did not differ between groups. These seemingly contradicting results relate to the difference in stimulus quality; while Randall-Selitto measures static hyperalgesia generally confined to the area of primary hypersensitivity, fine von Frey filaments can detect punctuate mechanical hypersensitivity expanding beyond the primary site of hypersensitivity (Jensen & Finnerup, 2014). Overall, the results of this study suggest that CIBP is associated with the development of mechanical hypersensitivity at distant sites, as has been previously described in mice (Guedon *et al.*, 2016) and humans (Scott *et al.*, 2011). On the other hand, we did not observe any differences in heat pain thresholds between sham and cancer animals, suggesting thermal sensitivity is intact in mice with a moderate pain phenotype, even though others have previously reported a reduction in paw withdrawal thresholds to radiant heat (Hargreaves) in murine models of CIBP (Gu *et al.*, 2009; Abdelaziz, Stone & Komarova, 2015).

### 3.3 Increased sensitivity of cutaneous afferents innervating the paw in animals with CIBP

Several lines of evidence in the current study point towards changes in excitability at the level of the DRG, which contributes to the peripheral drive of nociceptive signaling in CIBP. We found that a higher proportion of L3 cutaneous plantar afferents respond to pinch, as opposed to heat, in cancer bearing animals. This is contrary to what we observed in sham animals and reflects the behavioural phenotype of mechanical hypersensitivity associated with CIBP in our model. Similar changes in excitability of cutaneous afferents have been reported in a model of osteoarthritis associated with secondary hypersensitivity, with afferents innervating the hind paw showing increased responses to pinch as revealed by *in vivo* calcium imaging (Miller *et al.*, 2018). Our data also shows a significantly higher response intensity of mechanoresponsive plantar afferents in cancer bearing animals, which further supports the data demonstrating that mechanosensitive cutaneous neurons are sensitized in CIBP. Similarly, *in vivo* electrophysiological recordings from L4 DRG soma have also found that C-Aβ- and Aδ-fibers become more excitable, with decreased mechanical thresholds in rats with CIBP (Zhu *et al.*, 2016). A more recent study of CIBP in rats reported a recruitment of silent small-to-medium sized muscle afferents surrounding the cancer bearing bone in response to knee compression (Kucharczyk *et al.*, 2020).

We observed a substantial increase in polymodal responses of cutaneous afferents in cancer bearing animals, which may reflect a phenotypic shift in thermo-specific DRG neurons that acquire sensitivity to mechanical stimuli. This may underlie the observed increase in response intensity of heat-responding nociceptive neurons in cancer bearing animals, although this was not sufficient to induce behavioural changes in noxious heat thresholds. In line with this, a previous study reported that subcutaneous administration of inflammatory soup produces a response to mechanical stimulation in previously silent mechanically insensitive heat nociceptors (Smith-Edwards *et al.*, 2016). Moreover, we have also previously reported that intraplantar Prostaglandin E_2_ leads to a substantial increase in polymodal responses in mice and encourages a phenotypic switch in response modalities of DRG neurons (Emery *et al.*, 2016).

We observed a trend of increased DRG coupling in response to mechanical, but not thermal stimuli, although size distribution of these recruited cells did not differ between sham and cancer animals. A substantial increase in neuronal coupling events has previously been reported for both inflammatory and neuropathic pain, with both imaging and behavioural data indicating this phenomenon is more prominent for mechanical, rather than thermal hypersensitivity (Kim *et al.*, 2016). The discrepancy in effect size between these findings and the current study may depend on our approach to image at the level of L3, whereas L4 DRG contain somas of the majority of neurons with receptive fields in the hind paw (da Silva Serra *et al.*, 2016). While CIBP shares some features of inflammatory and neuropathic pain, it is seen as a distinct pain state (Middlemiss, Laird & Fallon, 2011; Falk & Dickenson, 2014) for which cross-activation of sensory neurons may play a minor role in peripheral sensitization.

### 3.4 Response properties of bone afferents are unaffected in mice with moderate bone cancer pain

To investigate the response properties of bone afferents to acute stimulation, a 10μl solution was injected in the mouse femur and activity of DRG neurons was recorded by *in vivo* calcium imaging. The time required to inject a 10μl solution in the intramedullary cavity of the mouse femur was significantly increased in animals with CIBP and the pressure of the injected solution tended to be higher in cancer bearing animals. Overall these findings indicate that an enhanced intraosseous pressure is created by the presence of tumour (Sottnik *et al.*, 2015). To stimulate bone afferents, a solution of saline was injected to produce increased intraosseous pressure (Nencini & Ivanusic, 2017). Citric acid injection was used to simulate the acidic microenvironment, which is thought to contribute to sensitization of primary afferents innervating cancerous tissue (Yoneda *et al.*, 2011; Yoneda, Hiasa, Nagata, Okui & F. White, 2015; Yoneda, Hiasa, Nagata, Okui & F.A. White, 2015) In both cases, we were surprised to find that unlike primary afferents innervating the glabrous hindpaw skin, the response properties of bone afferents were unaffected by the presence of cancer. This is in line with a recent study indicating that muscle afferents innervating the tissue surrounding the cancer bearing bone contribute to peripheral sensitization in rats, rather than bone innervating neurons themselves (Kucharczyk *et al.*, 2020). Similarly, electrophysiological studies point towards ectopic activity in cutaneous C-fibers surrounding the cancer bearing bone (Cain *et al.*, 2001). Overall these results suggest bone afferents are not mediating nociceptive processing in intermediate to late stage cancer. However, we cannot exclude the potential role of bone marrow afferents in early establishment of CIBP, as during disease progression the fibers innervating the bone undergo a continuous cycle of sprouting, degeneration and resprouting (Peters *et al.*, 2005; Jimenez-Andrade *et al.*, 2011)

### 3.5 Conclusion

We observed that the majority of afferents innervating the mouse femoral bone marrow express Na_v_1.8, and that the response properties of these DRG neurons to both pressure and acidic stimuli are unaffected in a mouse model of CIBP. However, cancer bearing animals did exhibit increased intraosseous pressure, as evidenced by prolonged times needed for the injection of a solution into tumour-bearing femurs. Surprisingly, we found that cutaneous afferents innervating the glabrous skin of the hindpaw in animals with CIBP show increased excitability at the level of the soma, as well as a phenotypic shift from thermosensitivity to increased mechanosensitivity. These findings are consistent with the behavioural phenotype of secondary mechanical hypersensitivity observed in cancer bearing mice compared to sham controls. Our data supports previous findings that pain behaviour and disease progression in a mouse model of CIBP occurs independently of bone marrow-innervating afferents, and is likely driven by an increase in polymodal neurons innervating secondary sites.

## Acknowledgments

This work was supported by the European Commission’s Horizon 2020 Research and Innovation Programme (Marie Skłodowska-Curie grant 642720), Versus Arthritis U.K., and the Wellcome Trust. We thank Edward C. Emery and other members of the Molecular Nociception Group for help and advice.

## Author Contributions

LdC, JNW and SS conceived the study, designed the experiments, interpreted the results. LdC performed the experiments with the help of APL and SSV, analyzed the data and wrote the manuscript. SS and JNW provided suggestions and contributed to writing the manuscript.

## 4 Materials and Methods

### 4.1 Cell culture

LL/2 Lewis Lung carcinoma cells (ATCC) were cultured in DMEM supplemented with 10% FBS and 1% Penicillin/Streptomycin for at least 2 weeks prior to surgery. Cells were split at 70-80% confluence two or one day prior to surgery (cell culture reagents supplied by Thermo Fisher). On the day of surgery cells were harvested with 0.05% Trypsin-EDTA, resuspended in DMEM at a final concentration of 4×10^7^ cells/ml and kept on ice till use. Cells were counted before and after intrafemoral injection to confirm viability.

### 4.2 Animals

For genetic labelling of neuronal subsets homozygous Rosa-flox-stop tdTomato mice (Madisen *et al.*, 2010) were crossed with homozygous Na_v_1.8-cre (Scn10a^tm2(cre)Jnw^) (Nassar *et al.*, 2004), Tmem233-cre (generated in our lab), or Pvalb-cre (B6;129P2-*Pvalb*^*tm1(cre)Arbr*^/J) (Hippenmeyer *et al.*, 2005) homozygous mice. *In vivo* calcium imaging experiments were performed using heterozygous Pirt-GCaMP3-expressing mice (at least 12 weeks old; male and female) on a C57BL/6J background, generated by X.D. (Johns Hopkins University, Baltimore, MD) (Kim *et al.*, 2014). Where indicated homozygous Pirt-GCaMP3 and Rosa-flox-stop tdTomato mice were crossed with homozygous Calb1-cre (Nigro, Hashikawa-Yamasaki & Rudy, 2018) or Tmem233-cre mice. For genotyping, genomic DNA was isolated from ear tissue. Primers used for PCR are summarized in Table S 1. Mice were housed in groups of 2-5 animals with a 12-hour light/dark cycle and allowed free access to water and standard diet. All animals were acclimatized for 2 weeks before the start of the experiment. All experiments were performed with approval of personal and project licenses from the United Kingdom Home Office according to guidelines set by the Animals (Scientific Procedures) Act 1986 Amendment Regulations 2012, as well as guidelines of the Committee for Research and Ethical Issues of IASP. Any mice that developed limping within 4 days after surgery were excluded (n=2). Cancer animals that failed to develop a reduction in limb use score over the course of 20 days (n =8) were also excluded.

### 4.3 Surgery

Cancer cells or DMEM were administered intrafemorally as previously described (Minett *et al.*, 2014; Bangash *et al.*, 2018), with a few adaptations to allow for injection of a retrograde tracer. Briefly, animals were anaesthetized with isoflurane. An incision was made in the skin above the patella. The patella and the lateral retinaculum tendons were loosened to move the patella to the side and expose the distal femoral epiphysis (Falk *et al.*, 2013). A 30-gauge needle was used to make a hole through the growth plate till the medullary cavity was reached. The needle was removed and a Hamilton syringe with an attached canulae was used to inject 5μl of 2×10^5^ LL/2 cells or DMEM medium (sham). This was kept in place for 2 min to allow cells to set. The canulae was quickly replaced with a new canulae to deliver 1 μl of 2% Fast Blue solution in water. Another 5 min were left to allow for the tracer to set. The hole was closed with bone wax (Johnson&Johnson) and the wound irrigated with sterile saline. The patella was moved back in place. The skin was sutured with 6-0 absorbable vicryl rapid (Ethicon), and Lidocane spray (Intubeaze, 20mg/ml, Dechra) applied to the wound. For colocalization experiments only Fast Blue was injected.

### 4.4 Behavioral tests

For behavioral experiments, animals were acclimatized to the equipment for at least 2 days prior to testing. The experimenter was blind to the groups. Limb use score and weight bearing was performed on all animals. For von Frey data some animals were excluded as withdrawal thresholds were measured before the animals reached a limb use score of 2 (see below). All other tests were performed on different subset of animals to accommodate for multiple testing and avoid stress evoked by multiple procedures carried out on a single day (detailed in Table S2).

#### 4.4.1 Limb use score

Mice were allowed to freely move around in a glass box (30×45 cm) for 10 min of acclimatization. Then each mouse was observed for a period of 4 min and the use of the affected limb was scored from 4 to 1 as follows. 4: Normal use of the limb; 3: slight limping, characterized by preferential use of the unaffected limb when rearing; 2: clear limping; 1: clear limping and partial lack of use of the limb. A limb score of 1 or 2 was used as endpoint for cancer animals.

#### 4.4.2 Weight bearing

Changes in weight-bearing were measured using an Incapacitance Meter (Linton Instrumentation) consisting of two scales. The mouse was allowed to place its head and upper body into a plastic tube to reduce stress and the hind limbs were positioned each on one of the scales. The load of each limb on the scale was measured for 5 sec in which the mouse was still. Measurements were taken in triplicate, changing the position of the hind legs after each trial. The average weight-bearing ratio was calculated as the weight placed on the affected limb divided by the total weight on both hind limbs.

#### 4.4.3 Mechanical sensitivity with von Frey

Mechanical hypersensitivity was measured using the up-down method to obtain the 50% withdrawal threshold (Chaplan *et al.*, 1994). In brief, mice were placed in darkened enclosures with wire mesh floor and left to habituate for at least 1 hr till movement was reduced to a minimum. Filaments were applied perpendicular to the plantar surface for 3 sec. Interval between stimuli was at least 30 sec. Starting from a 0.4 g filament, the response was recorded as negative for no reaction or as positive for paw withdrawal. A positive response resulted in a decrease in filament strength for the next stimulation, a negative response in increased strength. To determine the optimal threshold six responses in proximity of the 50% threshold are required. Thus, starting from the point at which the response to a filament changed from positive to negative or negative to positive, five further responses were recorded. When continuous positive responses were observed to the minimum stimulus of 0.008 g this set cut-off was used as the 50% withdrawal threshold. In the other cases, the pattern of responses obtained was used to calculate the 50% threshold = (10[χ+κδ])/10,000), where χ is the log of the final von Frey filament used, κ the tabular value for the pattern of responses, and δ the mean difference between filaments (in log units).

#### 4.4.4 Non-noxious mechanical sensitivity with palpation

Palpation was performed as previously described (Schwei *et al.*, 1999). Animals were gently restrained and their affected paw was held in place, while non-noxious pressure was applied by holding the distal femur between thumb and index every 1 sec for a total period of 2 min. The animal was then transferred into a darkened enclosure with glass flooring and nocifensive responses, such as guarding, licking, and lifting of the affected limb were quantified for a period of 2 min.

#### 4.4.5 Noxious punctate mechanical sensitivity with Randall-Selitto

The previously described Randall-Selitto apparatus was used (Takesue, Schaefer & Jukniewicz, 1969). To determine mechanical pain thresholds of the paw, animals were restrained by holding the scruff of the neck, and increasing pressure applied till a behavioural response, such as struggling, withdrawal of the paw, or vocalisation were observed. The test was repeated 3 times for each animal.

#### 4.4.6 Noxious heat sensitivity with hot-plate

The mouse was placed into the hot-plate apparatus (Ugo Basile), which was held at a temperature of 50°C. The test ended when the animal showed a withdrawal response or licked one of the hind paws. Cut off time was set to 45s.

#### 4.4.7 Immunohistochemistry

Mice were terminally anaesthetized 5-7 days after injection of Fast Blue with an intraperitoneal (i.p.) injection of sodium pentobarbitone (200 mg/kg) (Pentoject^®^, AnimalCare). Mice were perfused with ice-cold heparinized saline (10 U/ml heparin in 0.9% w/v NaCl), followed by 4% paraformaldehyde (PFA) (Sigma-Aldrich) solution in 0.1 M phosphate buffer (PB) (pH=7.4). DRGs were post-fixed in the same fixative solution for 2 hr at 4°C, embedded in OCT compound (Tissue-Tek^®^, Sakura) and left to set on dry ice, then stored at −80°C till sectioning. L3 DRG were serially sectioned at 80 μm (n=3, Na_v_1.8 tdTomato, n=2 Pvalb tdTomato) or 11 μm thickness and collected on electrostatically charged slides (Superfrost^®^ Plus, Thermo-Scientific). Slides were left to dry at RT and then stored at −80 °C. Tissues were removed from −80 °C, left to acclimatize to RT and then washed 3×5min in PBST (0.3% Triton X in 0.1M PBS). To reduce background signal slides were blocked for 1 hr in 1% bovine serum albumin (BSA, Sigma-Aldrich) in PBST, followed by 3×5min washes in PBST. Incubation with primary antibody in blocking buffer was performed ON at RT for TrkA (R&D AF1056, 1:1000). The next day, slides were washed 3×5 min in PBST and incubated with secondary antibodies (chicken anti-goat IgG Alexa Fluor 488 (ThermoFisher: A-21467, 1:1000) in blocking buffer for 2 hr at RT, followed by 3×5 min washes in PBS. Slides were dried in the dark at RT and mounted with Vectashiled Hardset Antifade mounting medium (Vector Laboratories) and coverslipped. Slides were either imaged directly or stored at −80°C.

### 4.5 *In vivo* calcium imaging

Mice were anesthetized using ketamine (120 mg/kg) (Fort Dodge Animal Health Ltd.) and medetomidine (1.2 mg/kg) (Orion Pharma). The depth of anaesthesia was assessed by pedal reflexes, breathing rate, and whisker movement. Throughout the experiment, the body temperature of the animal was kept at 37°C using a heated mat (VetTech). A dorsal laminectomy was performed at spinal level L2–L4 and the L3 DRG was exposed for imaging as previously described (Emery et al., 2016). Artificial spinal fluid (120 mM NaCl, 3 mM KCl, 1.1 mM CaCl2, 10 mM glucose, 0.6 mM NaH_2_PO_4_, 0.8 mM MgSO_4_, 18 mM NaHCO_3_ (pH 7.4) with NaOH) was constantly perfused over the exposed L3 DRG during the procedure to maintain tissue integrity. The lateromedial aspect of the left femur was exposed, by separating the biceps femoris posterior from the biceps femoris anterior. A hole was drilled at about 1 cm from the distal femur head, a canula was inserted and fixed in place with dental cement. In vivo imaging was performed using a Leica SP8 confocal microscope (Dry ×10, 0.4-N.A. objective with 2.2-mm working distance, Leica). Scans were taken at a bidirectional scan speed of 400-600Hz at a resolution of 512 × 512 pixels in either one or two z-planes. Pinhole A.U. was kept between 1.22 and 3.85 to visualize single cells accurately. A laser line of 405 nm, 488 nm were used to excite Fast Blue and GCaMP3 respectively. The collection of the resulting emission was system optimized to maximize yield and minimize crosstalk (Leica Dye Finder, LASX software; Leica).

#### 4.5.1 Peripheral stimulation

Study design of peripheral stimulation is shown in Figure 1. *In vivo* calcium imaging of L3 DRG was performed once cancer bearing animals reached limb score 2, representing a time point for significant sensory and motor dysfunction. (1) First, the glabrous skin of the ipsilateral hind paw was stimulated in order to activate calcium transients in cutaneous plantar afferents. Tweezers were used to apply pressure (pinch stimulation) for 3 sec across the dermatome covering L3-L4 for mechanical stimulation, followed by transient immersion of the paw in hot water (55 °C) and ice-cold water (0 °C) for 10 sec for thermal stimulation. Each stimulus application was separated by 30 sec. (2) Second, for activation of bone marrow afferents a 10 μl solution of either saline (0.9% NaCl) or 0.1 M citric acid (pH=4), containing 2.5 mg/ml Blue Evans was injected in the mouse femur. Bone marrow afferents were counted as responders if there was a change in unevoked baseline fluorescence either during the injection or within 30 sec after the injection. In a subset of animals the solution was delivered by a pump-controlled system (Harvard Apparatus Plus), where a 1 mm syringe was connected to both pressure gauge and a canula containing 40 μl volume. The gauge recorded changes in pressure with a maximum limit reading of 250 mmHg during the constant flow rate set at 10 ul/sec until at total of 10 μl solution was delivered. All but two animals reached the maximum limit reading before 10 μl solution was injected, and we therefore assumed that pressure was still rising above this value. In another subset of animals, pressure was applied manually using a syringe in order to produce faster changes in intraosseous pressure. (3) Finally, cutaneous plantar afferents were stimulated again as described in point 1. After *in vivo* calcium imaging a z-stack was recorded for counting the overlap between Fast Blue+ and tdTomato labelled cells. A necropsy was performed to confirm that the injected solution stayed within the femur and confirm that only neurons in the ipsilateral DRGs were labelled with retrograde tracer Fast Blue.

#### 4.5.2 Image analysis

All *in vivo* imaging data was acquired with the LAS-X analysis software (Leica) and analysed with ImageJ. All images were stabilized for XY movement using the TurboReg plug-in (Thévenaz, Ruttimann and Unser, 1998), with all images being registered to a stable image of the series. Raw traces of calcium signals were generated through the free hand selection tool of regions of interest (ROIs) surrounding cell bodies which responded to stimulus application. Area was used to determine average cell size of responding cells and average pixel intensity as a measure of change in calcium transients. Data was analysed by a combination of Matlab R2017a and Microsoft Office Excel 2013. Raw traces were first smoothed by averaging the preceding 4 frames of any test frame to reduce noise. To determine if a neuron was responsive to a given stimulus, the derivative of each frame was taken as ΔF/Δt. Neurons were counted as responders to a given stimulus if 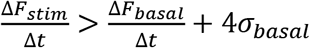, where F_stim_ is the maximum derivative value within a given window of stimulus application, F_basal_ is the average of derivative values in a 10sec time window preceding stimulus application, and σ_basal_ is the SD of the baseline derivative values. All neurons identified as responders were double-checked visually to avoid signal contamination by cells with partially overlapping ROIs. To generate normalised data for each trace, the following equation was applied 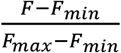. Coupled responses were determined manually and defined as two or more cell bodies within 5μm of each other responding to the same stimulus application. Relative presence of coupling events was determined by dividing the number of coupled responses by the total number of responses to that particular stimulus.

### 4.6 Statistical analysis

Statistical analysis was performed using GraphPad Prism 8. Log-rank test was used to compare survival distributions between groups. Welch’s t-test (two-tailed) and one-way ANOVA were used to compare means between two or more groups, respectively. Kolmogorov-Smirnov test was used to compare difference between two distributions. Comparison of multiple timepoints between groups was performed by two-way ANOVA with Bonferroni post-hoc test. Data is presented as mean ± standard error of the mean (S.E.M.), and significance as: *p<0.05; **p<0.01; ***p<0.001, ****p<0.0001.

---

**Figure S 1.**
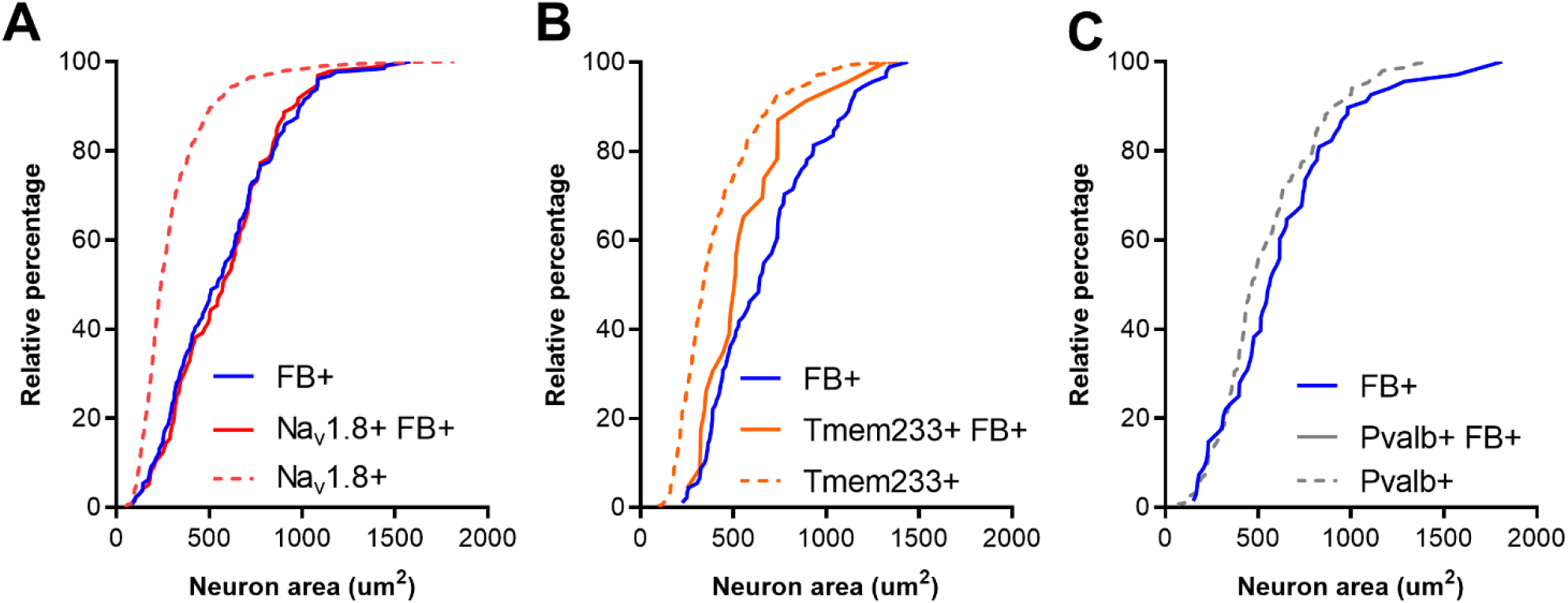
Size distribution of retrogradely labelled femoral bone marrow afferents within each genetically labelled neuronal subset. (A) Size distribution of the whole Na_v_1.8-cre+ population (ticked red line, n=839), all retrogradely labelled bone afferents (blue line, n=129), and those expressing Na_v_1.8 (red line, n=97). (B) Size distribution of Tmem233-cre+ population (ticked orange line, n=290), all retrogradely labelled bone afferents (blue line, n=91), and those expressing Tmem233 (orange line, n=22). (C Size distribution of Pvalb-cre+ population (ticked grey line, n=151), all retrogradely labelled bone afferents (blue line, n=68), and those expressing Pvalb (no line depicted, as n=1).

**Table S 1.**
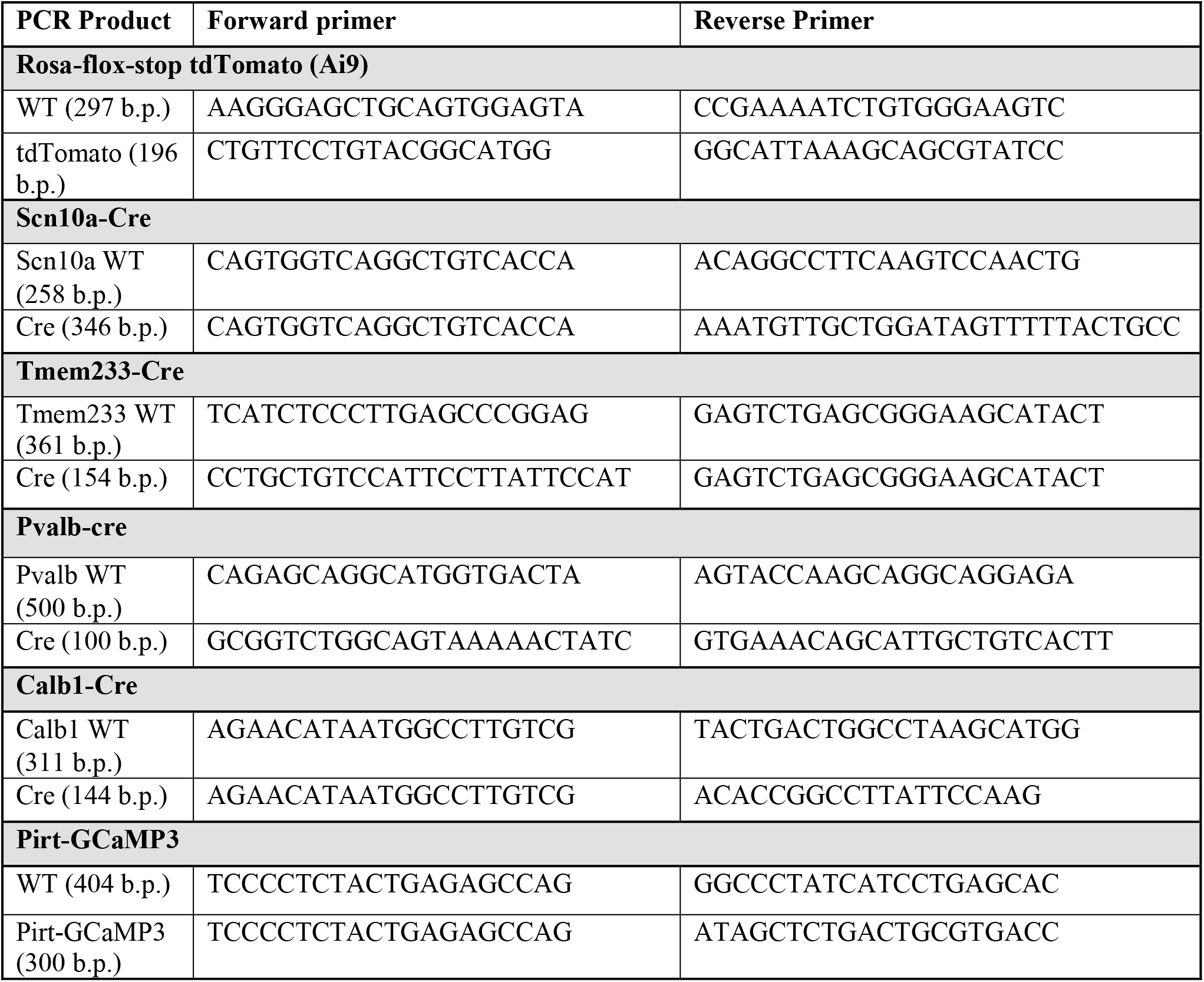
Primers used for genotyping.

**Table S 2.** Overview of animals used for behavioural testing.

| Experiment | Figure | Number of animals | Strain |
| --- | --- | --- | --- |
| Limb use score, weight bearing | Figure 2A-B | Sham: n=22<br>Cancer: n=17 | Sham: Pirt-GCaMP3 (n=11), Pirt-GCaMP3/Calb1-cre<sup>tdTomato</sup> (n=3), Pirt-GCaMP3/Tmem233-cre<sup>tdTomato</sup> (n=8)<br>Cancer: Pirt-GCaMP3 (n=12), Pirt-GCaMP3/Calb1-cre<sup>tdTomato</sup> (n=1), Pirt-GCaMP3/Tmem233-cre<sup>tdTomato</sup> (n=4) |
| von Frey | Figure 2C | Sham: n=21<br>Cancer: n=12 | Same as Limb use score and weight bearing, but excluding the following animals as behavioural testing at the endpoint was not possible:<br>Cancer: Pirt-GCaMP3 (n=4), Pirt-GCaMP3/Tmem233-cre<sup>tdTomato</sup> (n=1)<br>Sham: Pirt-GCaMP3 (n=1) |
| Noxious palpation | Figure 2D | Sham: n=11<br>Cancer: n=5 | Sham: Pirt-GCaMP3/Calb1-cre<sup>tdTomato</sup> (n=3), Pirt-GCaMP3/Tmem233-cre<sup>tdTomato</sup> (n=8)<br>Cancer: Pirt-GCaMP3/Calb1-cre<sup>tdTomato</sup> (n=1), Pirt-GCaMP3/Tmem233-cre<sup>tdTomato</sup> (n=4) |
| Randall-Selitto | Figure 2E | Sham: n=7<br>Cancer n=6 | Pirt-GCaMP3 |
| Hot-plate | Figure 2F | Sham: n=4<br>Cancer: n=6 | Pirt-GCaMP3 |

